# An ocean of opsins

**DOI:** 10.1101/2024.11.07.622103

**Authors:** Giacinto De Vivo, Eric Pelettier, Roberto Feuda, Salvatore D’Aniello

## Abstract

In this study, we explored the diversity and evolution of opsins using meta-omic data from the TARA Oceans and TARA Polar Circle expeditions, one of the largest marine datasets available. By using molecular clustering and phylogenetic analyses, we identified and characterized opsins across various taxonomic groups and gene subfamilies. Our results indicate that the majority of opsin sequences belong to arthropods and vertebrates. However, we also detected sequences from all major opsin subfamilies, including r-opsin, c-opsin, xenopsin, and Group-4 opsins. Despite the broad taxonomic scope, no novel opsin gene families were discovered, suggesting a comprehensive understanding of opsin diversity in marine ecosystems. Additionally, we present novel opsin sequences from less-studied taxa, which may serve as a valuable resource for future research into opsin function and diversity.

## Introduction

Plankton forms the foundation of the marine food web and harbors a vast portion of the biodiversity of the oceans. To better understand this diversity, the TARA Oceans expeditions embarked on a global journey, collecting and characterizing plankton from ecological, morphological, and molecular perspectives (Karsenti et al., 2011; Bork et al., 2015; Pesant et al., 2015; de Vargas et al., 2015; Carradec et al., 2018; Salazar et al., 2019; Sunagawa et al., 2020). With over 210 sampling locations worldwide and depths reaching up to 1000 meters, TARA have generated one of the most taxonomically rich genomic and transcriptomic datasets available. This remarkable resource provides an unprecedented opportunity to explore the evolution of gene families across a wide array of marine organisms.

Despite extensive research into plankton diversity, much of the marine biome remains unexplored at the molecular level, especially in terms of gene families crucial for ecological adaptations. Among these gene families, opsins hold a particularly important role for species interactions and ecology. They are key proteins required for animal vision, acting as light sensitive molecules that enable photoreception (Terakita, 2005; Porter et al., 2012). Traditionally opsins have been categorized in three main paralogue groups: ciliary opsin (c-opsin), rhabdomeric opsin (r-opsin), and Group-4 opsins (Go-opsins, neuropsin, RGR, and peropsin/retichronome) (Feuda et al., 2012; Fleming et al., 2020). However, recent studies have challenged this view, suggesting the existence of additional paralog groups. Indeed, new subfamilies such as xenopsin, non-canonical r-opsin, bathyopsin, chaopsin and pseudospin have been proposed (D’Aniello et al., 2015; Ramirez et al., 2016; De Vivo et al., 2023). The relationships between these subfamilies are still subject to debate. Some phylogenies suggesting the monophyly of Group-4 opsins and c-opsins with r-opsins sister group to them (Davies et al., 2010; Feuda et al., 2012; Feuda et al., 2014; Yoshida et al., 2015; Vöcking et al., 2017; Rawlinson et al., 2019; Bonadè et al., 2020; Fleming et al., 2020; McCulloch et al., 2023), while others support the monophyly of Group-4 and r-opsins (Porter et al., 2012). In contrast, xenopsins have been variably positioned, sometimes closer to c-opsins (Yoshida et al., 2015; Vöcking et al., 2017; Vöcking et al., 2021; McCulloch et al., 2023) and at other times closer to Group-4 opsins (Ramirez et al., 2016; Rawlinson et al., 2019; Bonadè et al., 2020).

A key limitation in understanding opsin diversity and evolution stems from a focus on well-characterized species, whose genomes or transcriptomes have been sequenced. This narrow taxonomic scope may obscure the full range of opsin diversity and hamper a comprehensive understanding of their evolutionary history. To address these gaps, this study investigates opsin diversity within the meta-transcriptomic (metaT) and meta-genomic (metaG) datasets of the TARA Oceans and TARA Arctic expeditions—the largest -omics dataset of marine plankton (Karsenti et al., 2011; Bork et al., 2015; Pesant et al., 2015; de Vargas et al., 2015; Carradec et al., 2018; Salazar et al., 2019; Sunagawa et al., 2020). This dataset offers an unprecedented chance to examine opsin diversity across a taxonomically and ecologically diverse range of marine organisms, providing new insights into the evolution and function of this critical gene family. We provide a quantitative description of opsin diversity, examining their taxonomic distribution and subfamilies composition within the TARA datasets, contributing to a better understanding of opsin evolution across the marine biome.

## Results

We analysed over 158 million eukaryotic gene sequences obtained from plankton samples across more than 210 sampling stations globally. Using a combination of BLAST, clustering, and annotation methods (Figure S1), we identified 2093 opsin sequences. To ensure a robust taxonomic classification, we integrated these sequences with 567 well-annotated reference sequences (2093 TARA sequences + 567 reference sequences; File S1). A phylogenetic reconstruction was then performed, using Maximum Likelihood under the WAG+CAT model on FastTree (Price et al., 2010) allowing us to map the diversity of opsins across various phyla.

Our results indicate that the vast majority of opsin (∼80%) were assigned to arthropods, which dominate zooplankton communities. Other taxa identified include cnidarians, acoelomates, molluscs, annelids, chaetognaths, rotifers, echinoderms, and vertebrates (Fig. 1A). The sequence diversity spanned across all r-opsin, c-opsin, xenopsin and Group-4 clades, with the exception for RGR opsins that have not been found (Fig. 1B). This absence of RGR opsins raises questions about potential lineage-specific losses or methodological biases, warranting further exploration. Among arthropods. we identified 1570 r-opsin, 189 c-opsin, 88 peropsin, 5 Go-opsin, and 36 neuropsins. Vertebrates, the second most represented group with a total of 110 opsin sequences, comprised 94 c-opsins and 16 peropsins. Rotifers were the third most abundant group, with 41 sequences in total: 5 r-opsins, 27 xenopsins, and 9 peropsins. Cnidarians, the fourth most represented group, had 13 sequences, all of which clustered with known cnidarian c-opsins. In chaetognaths, we found 2 xenopsins and 9 peropsins consistent with known absence of canonical visual opsins (e.g., r- and c-opsin) in this taxon despite having eyes (Wollesen et al., 2023). All 11 annelid opsins, as well as the single opsin found in acoelomates, belonged to the r-opsin clade. Among the molluscs, we identified 5 sequences: 2 r-opsins, 2 Go-opsins, and 1 neuropsin. The only echinoderm sequence in our dataset clustered with bathyopsins. Additionally, 9 tunicate sequences clustered ambiguously with cnidarian c-opsins, likely due to their high sequence divergence. However, based on reciprocal BLAST results, we classified them as peropsins/retinochromes. Finally, three additional sequences could not be classified with confidence and were discarded from further analysis.

**Figure 1.**
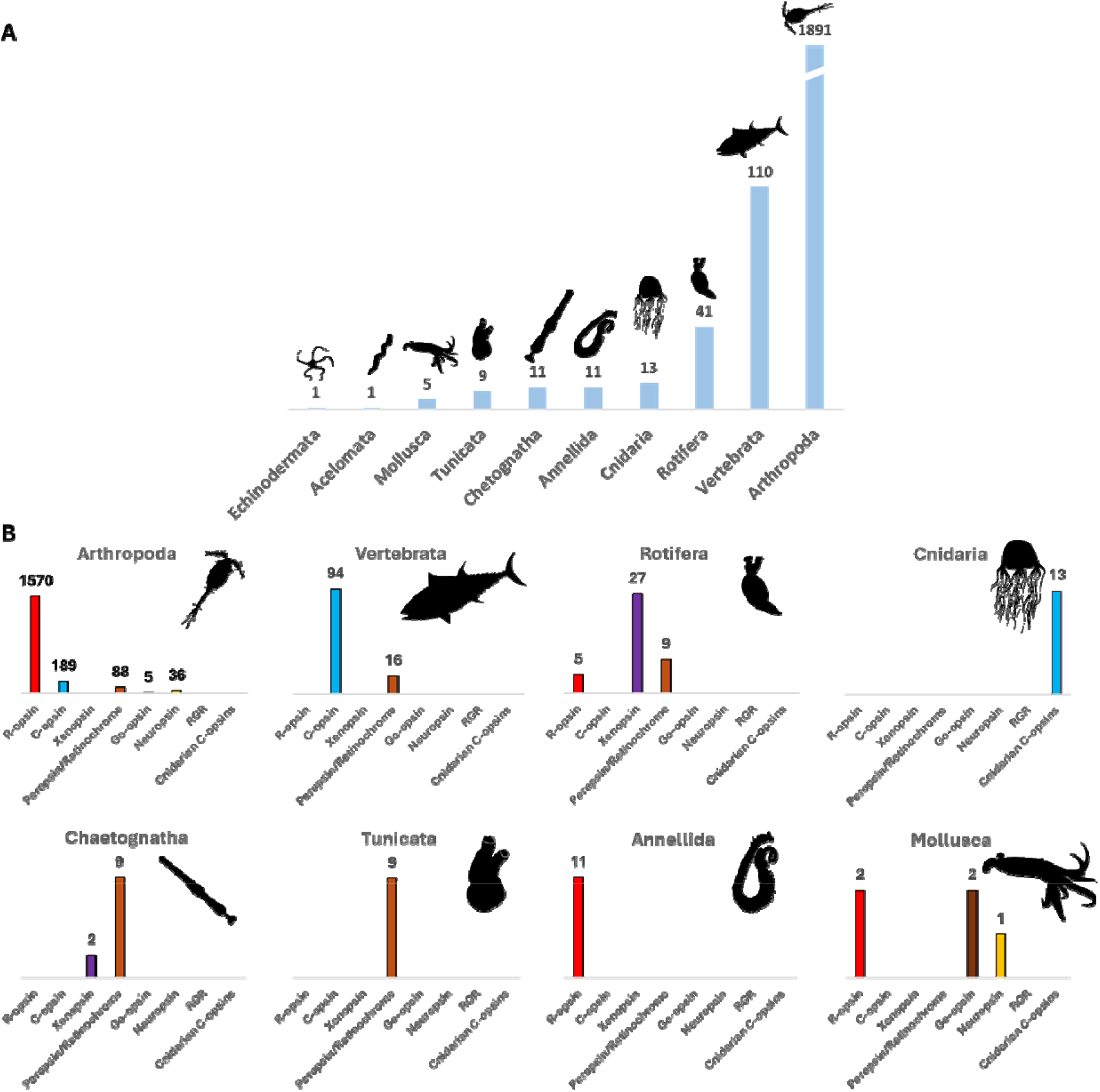
Opsin taxonomical distribution mined in TARA. (A) A plot showing the estimated number of opsin genes in TARA per taxonomical group. (B) Distribution of opsin subfamilies within different taxonomical groups. Animal silhouettes from Phylopic.

We then investigated the relationships among opsin subfamilies by performing a second phylogenetic analysis, aimed at reducing potential bias caused by the high number of arthropod sequences. To achieve this, we generated a second dataset in which 95% of arthropod sequences were removed (Dataset 2 = 298 TARA sequences + 567 reference sequences; File S2) and recompute the analysis using Maximum Likelihood under the WAG+CAT model on FastTree. Nodal support was estimated using the Shimodaira-Hasegawa test on the three alternate topologies (Price et al., 2010).

The resulting phylogenetic tree (Fig. 2 and Fig. S2) supports the existence of the four classical monophyletic groups: r-opsin with a bootstrap (BS) support of 0.99, c-opsin (BS=0.92), xenopsin (BS=0.97), Group-4 opsin (BS=0.94). Group-4 includes peropsin/retinochrome (BS=0.98), RGR (BS=0.99), Go-opsins (BS=0.85), and neuropsins (BS=0.97). The earliest emerging opsin group contains divergent sequences belonging to cnidarians, with r-opsin acting as the sister group to all other opsins. The second diverging group is c-opsin although this node is not well supported (BS=0.38) (Fig. 2; Fig. S3). Xenopsin, formed a clade with a group of cnidarian sequences that usually cluster with c-opsin (referred ‘cnidarian c-opsin’, BS=0.96) and all together with Group-4 they form a large clade (BS=0.80). A small group referred as bathyopsin (BS=0.86; Ramirez et al., 2016) is found as sister of the xenopsin/Group-4 clade (BS=0.92).

**Figure 2.**
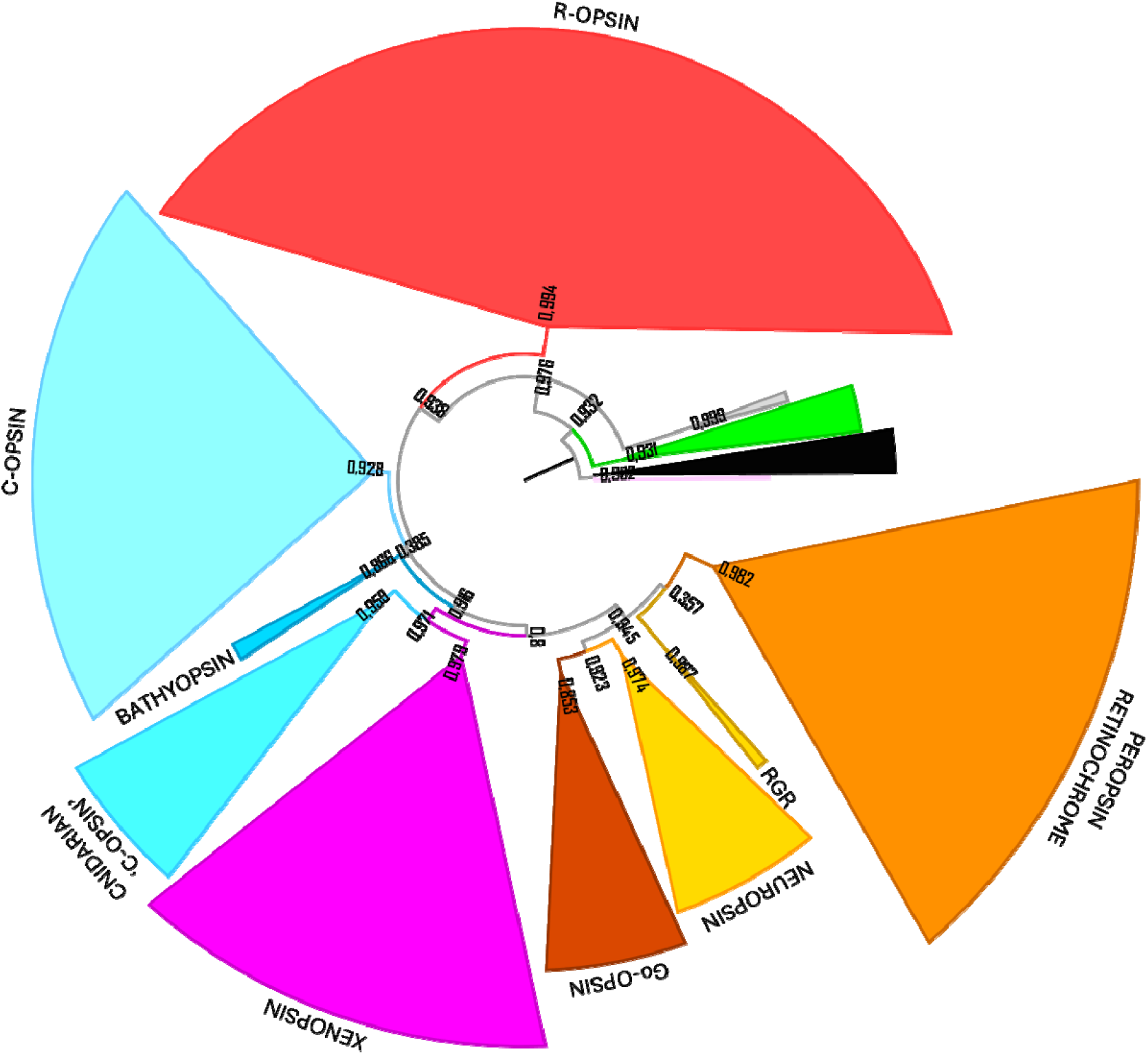
Maximum Likelihood phylogenetic tree (WAG+CAT) showing the relationships between the main opsin subfamilies: r-opsin (red), Group-4 opsin (orange, yellow, straw green, brown), xenopsin (magenta), c-opsin (cyan). Divergent cnidarian opsins are represented in grey, while the outgroups placopsin and pseudopsin are in pink and green, respectively. In this tree 95% of arthropod sequences have been removed. Bootstrap (B) values are indicated at the nodes.

## Discussion

Our results contribute to the study of metazoan opsin diversity and evolution and indicate that, despite mining 158 million unbiased eukaryotic gene sequences from plankton, we did not discover any new opsin gene subfamilies. This suggests that current knowledge of metazoan opsin diversity is approaching completeness, at least within the marine biome. However, the study highlights that this dataset, while extensive, may underrepresent certain opsin subfamilies.

Opsin subfamilies in our dataset appear to be unequally distributed. For instance, excluding the highly expressed visual opsins (c- and r-opsins), subfamilies such as peropsins and xenopsins are notably underrepresented. This could be linked to the extensive gene duplications observed in opsins involved in visual tasks, such as those that mediate colour vision. For instance, 77 out of 106 vertebrate c-opsins identified in this study belong to green-sensitive opsins of the rhodopsin-like 2 (Rh2) group, which have undergone substantial duplications in fishes (Hagen et al., 2023).

Moreover, our study supports the monophyly of c- and Group-4 opsins, consistent with previous research (Fleming et al., 2020; Feuda et al., 2015; Feuda et al., 2012), though certain evolutionary relationships, particularly for cnidarian and urochordate opsins, remain unresolved. This phylogenetic instability emphasizes the need for continued investigation into highly divergent opsin lineages and their evolutionary trajectories.

The functional roles of non-visual opsins, such as peropsins and retinochromes, also remain underexplored, despite their potential ecological importance. While the classification of major opsin subfamilies appears robust and well-supported, continued investigation into the functional roles and evolutionary histories of a large opsin diversity will be crucial to clarify their function as moonlighting protein (Feuda et al., 2022).

## Methods

### Data filtering

We initially mined opsin genes from a database containing 158 million gene sequences derived from meta-genomic and meta-transcriptomic samples collected during the TARA Oceans and TARA Arctic expeditions. Translated sequences from the TARA dataset, both genomic (SMAGs) and transcriptomic (MATOU-v1.5), were blasted (BLASTp) using Diamond against a precompiled opsin dataset representative on all the entire known opsin subfamilies as a query composed of 50 sequences. As a result, we obtained 14,176 putative opsin sequences were obtained (SMAGs: 5,266; MATOU: 8,910).

To filter out all non-opsin sequences that were collected by the initial BLAST search and to check if potential new opsin gene families were present, we proceeded through the following steps: 1) The sequences were merged with a larger opsin dataset consisting of 35,745 sequences obtained from UniProt and NCBI 2) Sequences shorter than 150 amino acids were removed from the dataset; 3) Only sequences with more than 4 and less than 9 transmembrane domains (TM), as predicted by SCAMPI (Peters at al., 2015), were retained for subsequent analysis. After these steps, a total of 43,228 sequences remained, of which 7,836 were from the TARA dataset.

To classify putative opsin in TARA dataset and given the computational limitation of performing a phylogenetic analysis using 43,228 sequences, we opted to split the dataset into subgroups based on protein similarity. The rationale behind this approach is that by grouping sequences with similar proteins, we should be able to identify all TARA sequences that exhibit some level of similarity to known opsins. To achieve this goal, we conducted an all-versus-all BLASTp in Diamond (e-value threshold: 1e^-10^). Sequences were then clustered based on their similarity using the Markov Cluster Algorithm (MCL) (van Dongen and Abreu-Goodger, 2019) with an inflation value set to 1.4. Each cluster was annotated using Eggnog v5.0 (Huerta-Cepas et al., 2019), and only the clusters that contained at least an annotated opsin were retained. This filtering process resulted in 369 clusters out of 735, of which 127 contained at least one opsin from TARA and 16 clusters were exclusively from TARA. This resulted in 2,317 putative opsins (1,801 from MATOU and 516 from SMAG). To confirm the identity of these 2,317 sequences, we BLASTP against the NCBI database. A total of 223 sequences were discarded, leaving a final set of 2,093 opsin sequences for subsequent analysis. The entire pipeline is outlined in Fig. S1.

### Phylogenetic analysis

For the phylogenetic analysis, we generated a combined dataset that included all 2,093 TARA opsin sequences and 567 previously annotated opsin sequences (Dataset 1; File S1). The 567 sequences included placopsins, pseudopsins, and melatonin receptors to serve as outgroup (Feuda et al, 2012; De Vivo et al, 2023; McElroy et al. 2024). The datasets were aligned using MAFFT-DASH v7.3 (server https://mafft.cbrc.jp/alignment/software/, option mafft --averagelinkage --reorder -- anysymbol --allowshift --unalignlevel 0.8 --dash --dashserver https://sysimm.ifrec.osaka-u.ac.jp/dash/REST1.0/ --maxiterate 0 --globalpair input) and trimmed using TrimAL with gap treshold of 0.1 (Capella-Gutiérrez et al., 2009; Rozewicki et al., 2019). Finally, phylogenetic analyses have been performed using FastTree (model WAG+CAT) (Price et al., 2010). Finally, taxonomic data were assigned by looking the tree topology and confirmed using reverse BLASTp on NCBI database.

To address the potential impact of the large number of similar sequences from closely related organisms—especially arthropods—on the phylogenetic results (e.g., node support values), we conducted a second phylogenetic analysis. In this analysis, we removed 95% of the TARA sequences assigned to arthropods from Dataset 1, resulting in Dataset 2 (File S2), and recomputed the phylogeny using the same methods as described above.

---

**Figure S1**

Bioinformatic pipeline was adopted to efficiently mine, cluster, and annotate opsins from TARA metaT (MATOU) and metaG (SMAGs) to be used for the phylogenetic analyses.

**Figure S2**

Phylogenetic analysis computed using all the 2093 sequences from TARA and an already annotated dataset of 567 reference sequences (i.e. Dataset 1).

**Figure S3**

Phylogenetic analysis computed using sequences from TARA where 95% of arthropod sequences were randomly subsampled (only 5% retained) and an already annotated dataset of 567 reference sequences (i.e. Dataset 2).

**File S1**

Alignment containing all the 2093 sequences from TARA and an already annotated dataset of 567 reference sequences (Dataset 1).

**File S2**

Alignment containing sequences from TARA where arthropod sequences were subsamples (only 5% retained) and an already annotated dataset of 567 reference sequences (Dataset 2).

## Data availability

Data are available on GitHub: https://github.com/Xodroont/Tara_Opsins MATOU-v1.5 and SMAGs data are available at http://www.genoscope.cns.fr/tara/

## Acknowledgements

We are grateful to Elijah Lowe, Luigi Caputi, Daniele Iudicone, Maria Ina Arnone for discussions on initial versions of the project. G.D.V. has been supported by a PhD fellowship funded by the Stazione Zoologica Anton Dohrn (Open University – Stazione Zoologica Anton Dohrn PhD Program). R.F. is supported by a University Research Fellowship (UF160226 and URF/R/221011) a Research Grant from the Royal Society (RGF\R1\181012).

Our survey was made possible by the sampling and sequencing efforts of the *Tara* Oceans Project. We are indebted to all who contributed to these efforts and other open-source bioinformatics tools for their commitment to transparency and openness. *Tara* Oceans (which includes the *Tara* Oceans and *Tara* Oceans Polar Circle expeditions) would not exist without the leadership of the *Tara* Oceans Foundation and the continuous support of 23 research institutes (https://oceans.taraexpeditions.org/). We also acknowledge the commitment of the CNRS and Genoscope/CEA. Some computations were performed thanks to the TGCC computing facility in France.

