## Supplementary figures and images for "An ocean of opsins"

### FigureS1

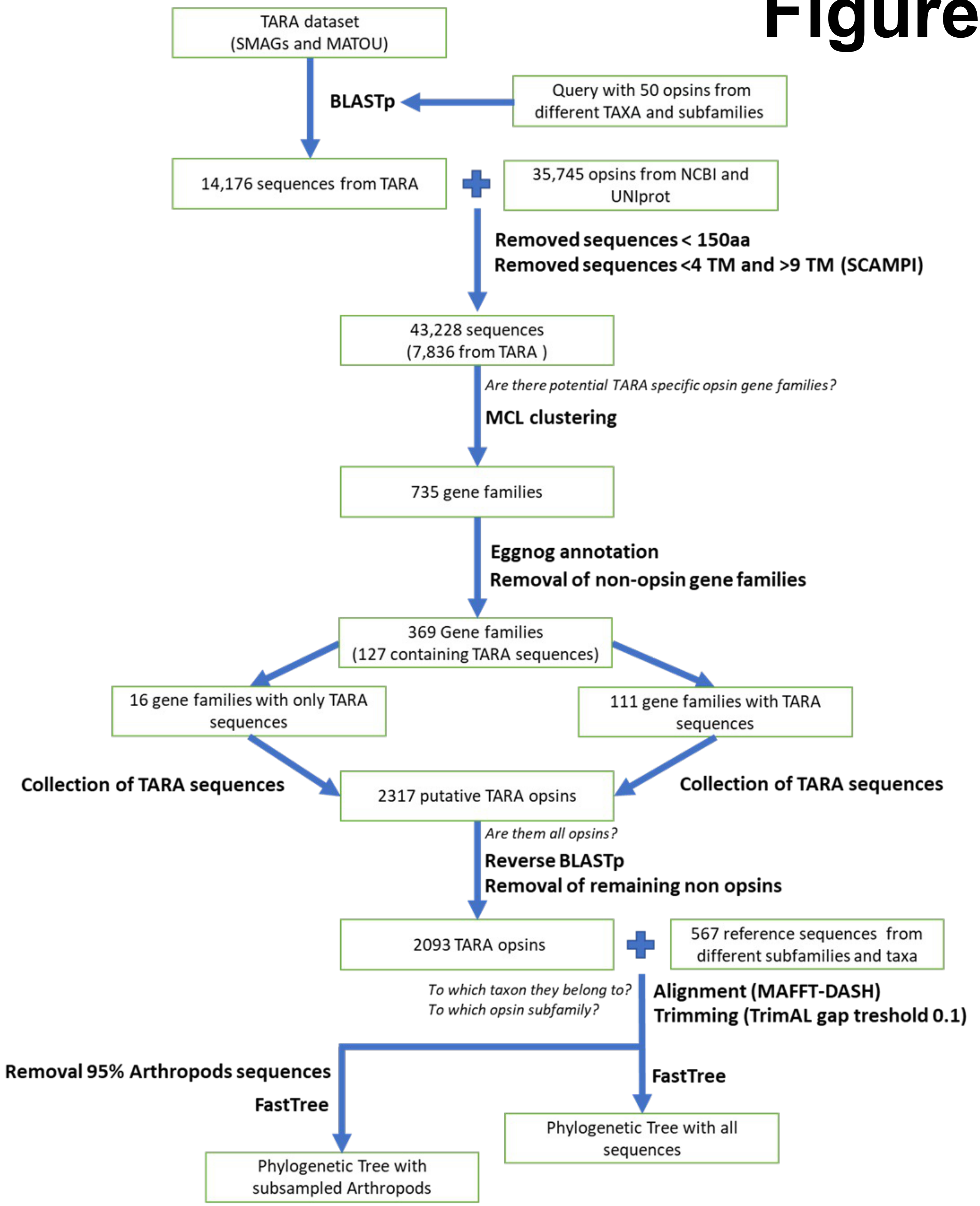

### FigureS3

Figure S3

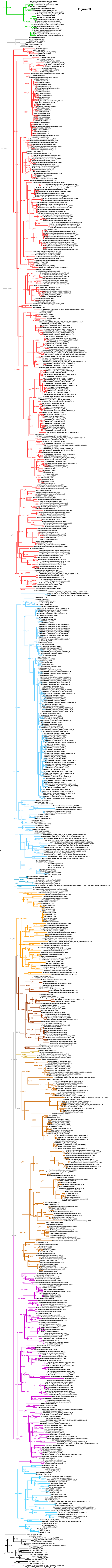
